# Longitudinal development of thalamocortical functional connectivity in 22q11.2 deletion syndrome

**DOI:** 10.1101/2023.06.22.546178

**Authors:** Charles H. Schleifer, Kathleen P. O’Hora, Maria Jalbrzikowski, Elizabeth Bondy, Leila Kushan-Wells, Amy Lin, Lucina Q. Uddin, Carrie E. Bearden

**Affiliations:** Department of Psychiatry and Biobehavioral Sciences, Semel Institute for Neuroscience and Human Behavior, University of California, Los Angeles, CA, USA; David Geffen School of Medicine, University of California, Los Angeles, CA, USA; Department of Psychiatry and Behavioral Sciences, Boston Children’s Hospital, Boston, MA, USA; Department of Psychiatry, Harvard Medical School, Boston, MA, USA; Department of Psychology, University of California, Los Angeles, CA, USA

**Keywords:** Copy Number Variant, Neurodevelopment, Psychosis, Autism, Thalamus, fMRI

## Abstract

**Background:** 22q11.2 Deletion Syndrome (22qDel) is a genetic Copy Number Variant (CNV) that strongly increases risk for schizophrenia and other neurodevelopmental disorders. Disrupted functional connectivity between the thalamus and somatomotor/frontoparietal cortex has been implicated in cross-sectional studies of 22qDel, idiopathic schizophrenia, and youth at clinical high risk (CHR) for psychosis. Here, we use a novel functional atlas approach to investigate longitudinal age-related changes in network-specific thalamocortical functional connectivity (TCC) in 22qDel and typically developing (TD) controls.

**Methods:** TCC was calculated for nine functional networks derived from resting-state functional magnetic resonance imaging (rs-fMRI) scans collected from n=65 22qDel participants (63.1% female) and n=69 demographically matched TD controls (49.3% female), ages 6 to 23 years. Analyses included 86 longitudinal follow-up scans. Non-linear age trajectories were characterized with general additive mixed models (GAMMs).

**Results:** In 22qDel, TCC in the frontoparietal network increases until approximately age 13, while somatomotor and cingulo-opercular TCC decrease from age 6 to 23. In contrast, no significant relationships between TCC and age were found in TD controls. Somatomotor connectivity in 22qDel is significantly higher than TD in childhood, but lower in late adolescence. Frontoparietal TCC shows the opposite pattern.

**Conclusions:** 22qDel is associated with aberrant development of functional network connectivity between the thalamus and cortex. Younger individuals with 22qDel have lower frontoparietal connectivity and higher somatomotor connectivity than controls, but this phenotype may normalize or partially reverse by early adulthood. Altered maturation of this circuitry may underlie elevated neuropsychiatric disease risk in this syndrome.

## Main Text

### Introduction

22q11.2 Deletion Syndrome (22qDel), also known as DiGeorge or Velocardiofacial syndrome (OMIM #188400, #192430), is a genetic disorder that occurs in approximately 1 in 4000 live births (1). This syndrome is one of the greatest genetic risk factors for schizophrenia, with at least 1 in 10 individuals with 22qDel having a comorbid psychotic disorder, and even higher rates after adolescence (2). Individuals with 22qDel also have high rates of autism and an increased incidence of intellectual disability (ID), attentional deficits, and anxiety disorders (3–5). 22qDel is caused by a copy number variant (CNV), consisting of a hemizygous deletion of 1.5-2.6 Mb of genetic material (∼46 protein-coding genes) from chromosome 22q (6). This provides a genetics-first framework for studying the biology underlying neurodevelopmental psychiatric disorders like schizophrenia (7,8).

Functional neuroimaging studies of psychosis spectrum disorders have consistently identified alterations in the functional connectivity (FC) of the thalamus, specifically, increased connectivity to somatomotor brain regions and decreased connectivity to frontoparietal associative regions compared to healthy controls (9–12). This marker is associated with conversion to psychosis in youth at clinical high risk (CHR) for the illness (13). In a cross-sectional study comparing individuals (ages 7-26) with 22qDel to typically developing (TD) controls, we observed a similar pattern of thalamic hyper-connectivity to the somatomotor network and hypo-connectivity to frontoparietal regions (14). This convergence of findings in idiopathic schizophrenia, CHR, and 22qDel may represent a shared phenotype relevant to psychosis risk. Interestingly, animal models of 22qDel implicate haploinsufficiency of the *Dgcr8* gene (deleted in 22qDel) in elevation of thalamic dopamine D2 receptors and age-related disruptions in thalamocortical synchrony (15,16) and may indicate an underlying neurobiological mechanism driving this dysfunction in 22qDel.

Thalamic dysconnectivity has additional cross-diagnostic relevance. Functional neuroimaging studies in autistic individuals have consistently observed altered thalamocortical FC (17–20). Many of these findings converge on increased connectivity within sensory networks. Furthermore, a broad convergence on disrupted thalamic connectivity has been identified in the functional connectomes from multiple idiopathic psychiatric conditions and neurodevelopmental CNVs (21,22).

The thalamus is a heterogeneous structure with dense reciprocal connections across the cortex (23). Structural and functional connectivity with the cortex across sensory and associative networks is thus a core organizational feature of the thalamus (24,25). Thalamocortical FC patterns specific to sensory and associative networks emerge early in development and have been identified in infancy (26). Recent studies of the relationship between age and thalamic FC have observed subtle developmental changes in sensory and associative network connectivity (27,28). Adolescence represents an important developmental window during which interactions between the thalamus and cortex have been hypothesized to shape prefrontal development, which may be disrupted in disorders like schizophrenia (29). One prior study of 22qDel has shown altered development of thalamic nuclei volumes, along with cross-sectional disruptions in FC (30).

In this longitudinal resting-state functional magnetic resonance imaging (rs-fMRI) study, we map age-related changes in thalamocortical FC in 22qDel and demographically matched TD control subjects, from childhood to early adulthood. To our knowledge, this is the first analysis of age-related changes in thalamic FC in 22qDel, and one of the first longitudinal analyses of thalamocortical FC in any population. Here, we use a novel functional atlas approach to compute network-specific thalamocortical functional connectivity (TCC) and generate non-linear mixed models of the relationship between age and TCC to test the prediction that development of frontoparietal and somatomotor TCC is altered in 22qDel. By examining the developmental trajectory of TCC in 22qDel, this study aims to shed light on the neurobiological mechanisms underlying the increased risk of psychosis and other neurodevelopmental disorders in this population.

### Methods

#### Participants

The total longitudinal sample consisted of 220 scans from 135 participants (6–23 years of age; 65 22qDel baseline; 69 TD controls baseline; see **Figure 1**), recruited from an ongoing longitudinal study at the University of California, Los Angeles (UCLA). 22qDel and TD participants were statistically matched based on baseline age, sex, dominant hand, fMRI movement (percent frames flagged based on displacement/intensity thresholds recommended by Power et al. 2012 (31)), as well as mean number of longitudinal visits and interval between visits, using appropriate tests (ANOVA or chi-squared; see **Table 1**). See **Supplemental Methods** for details on inclusion/exclusion criteria and clinical assessment procedures. After study procedures were fully explained, adult participants provided written consent, while participants under the age of 18 years provided written assent with the written consent of their parent or guardian. The UCLA Institutional Review Board approved all study procedures and informed consent documents.

**Figure 1.**
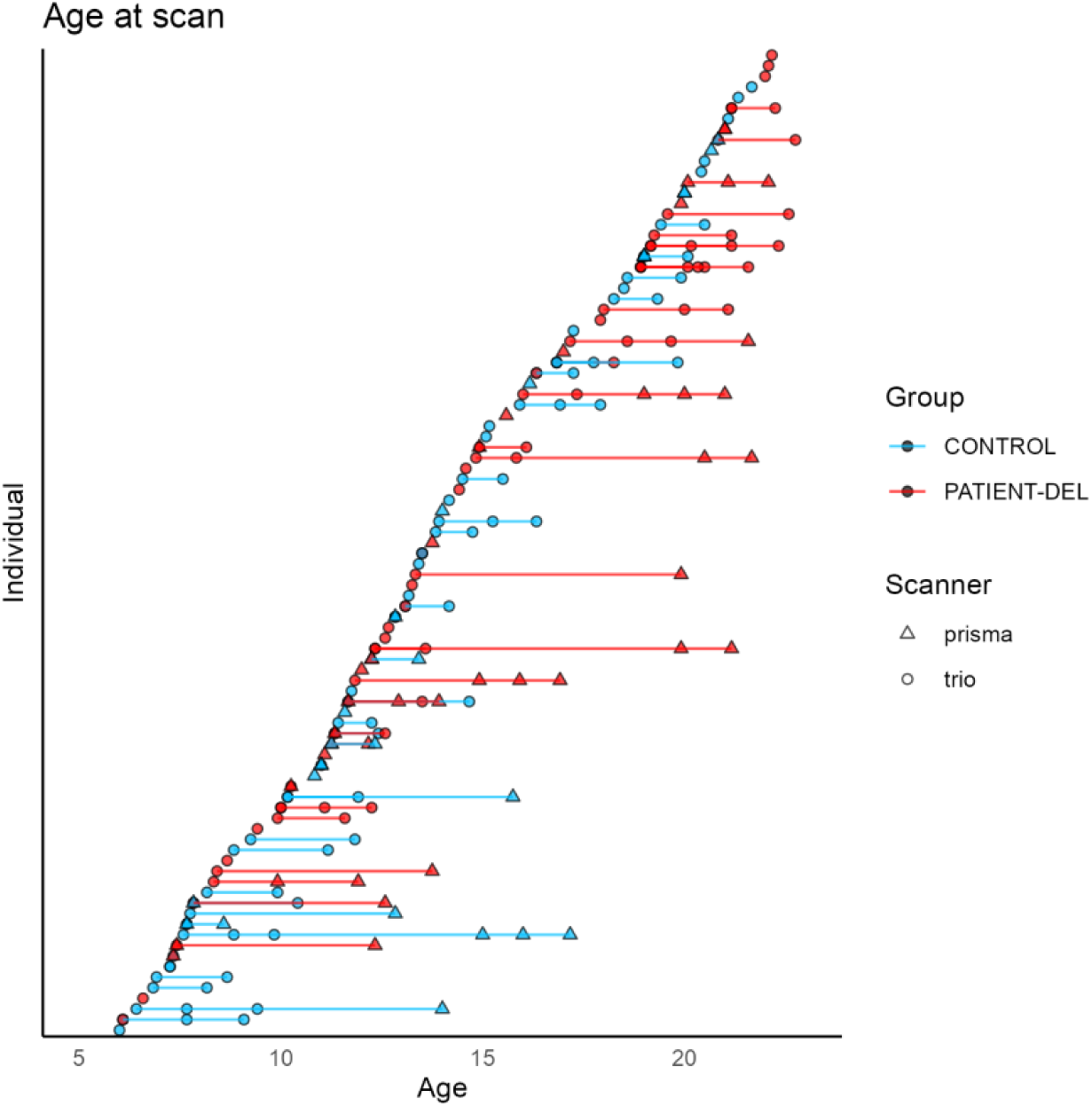
Participant age distribution. Typically developing controls in blue, 22qDel in red, with lines connecting follow-up visits from the same individual. Scanner type (Siemens Trio or Prisma) indicated by circle or triangle, respectively.

**Table 1.** Baseline demographics. 22qDel and TD controls with *p*-values for between group comparisons (based on ANOVA for continuous variables and chi-squared tests for categorical variables). Baseline cohorts are statistically matched based on age, sex, dominant hand, and fMRI movement, as well as mean number of longitudinal visits and interval between visits. Cognition was measured with the Wechsler Abbreviated Scale of Intelligence-2 (WASI-2). Prodromal (psychosis-risk) symptoms were assessed with the Structured Interview for Psychosis-Risk Syndromes (SIPS). Psychosis Risk Symptoms are operationalized here as having any score of 3 or greater (i.e., prodromal range) on any SIPS positive symptom item. Psychotic disorder diagnosis is based on structured clinical interview (SCID) for DSM-IV/V and includes schizophrenia, schizoaffective disorder, brief psychotic disorder, and psychotic disorder not otherwise specified.

|  | Control | 22qDel | *p*-value |
| --- | --- | --- | --- |
| n | 69 | 65 |  |
| Age, mean (SD) | 13.44 (4.76) | 14.39 (4.56) | 0.242 |
| Sex, n (%) Female | 34 (49.3) | 41 (63.1) | 0.151 |
| Handedness, n (%) Right | 31 (44.9) | 34 (52.3) | 0.561 |
| fMRI % movement, mean (SD) | 6.86 (9.93) | 9.84 (13.12) | 0.139 |
| WASI Full Scale IQ, mean (SD) | 112.15 (20.53) | 79.00 (12.70) | <0.001 |
| SIPS Positive total, mean (SD) | 1.18 (2.16) | 5.44 (5.61) | <0.001 |
| Psychosis Risk Symptoms, n (%) | 4 (5.8) | 20 (30.8) | 0.001 |
| Psychotic Disorder, n (%) | 0 (0.0) | 5 (7.7) | 0.058 |
| ADHD, n (%) | 5 (7.2) | 30 (46.2) | <0.001 |
| Autism, n (%) | 0 (0.0) | 36 (55.4) | <0.001 |
| Antipsychotic med, n (%) | 0 (0.0) | 5 (7.7) | 0.004 |
| Visit count, mean (SD) | 1.57 (0.85) | 1.72 (0.96) | 0.314 |
| Days between visits, mean (SD) | 653.21 (425.56) | 787.04 (549.92) | 0.296 |

#### Neuroimaging acquisition and processing

Rs-fMRI and high-resolution structural images were collected on two scanners (Siemens Trio and Siemens Prisma) at the UCLA Center for Cognitive Neuroscience, and processed with the Quantitative Neuroimaging Environment & Toolbox (QuNex) (32), which adapts the Human Connectome Project (HCP) preprocessing pipelines (33) for broader use. Additional processing of the fMRI time series included bandpass filtering, motion scrubbing for frames exceeding either a framewise displacement or signal change threshold (31), spatial smoothing, and regression of mean signal from ventricles, deep white matter, and mean gray matter (34). Scans with >50% frames flagged for motion were excluded. For a detailed description of preprocessing methods, see **Supplemental Methods** and previous work in a subset of these data (14).

For each scan, TCC was computed based on the correlation in fMRI signal between the thalamic and cortical components of nine networks (frontoparietal, somatomotor, cingulo-opercular, default mode, dorsal attention, auditory, posterior multimodal, primary visual, and secondary visual) defined by the Cole-Anticevic Brain-wide Network Partition, a recently developed whole-brain functional atlas (35) (see **Figure 2**). Data from the two scanners were harmonized using the longitudinal ComBat (longComBat) package in R (36), a linear mixed effects adaptation of the ComBat approach which uses empirical Bayes methods to estimate and remove site/batch effects with increased robustness to outliers in small samples compared with general linear model methods (37) (see **Supplemental Methods**).

**Figure 2.**
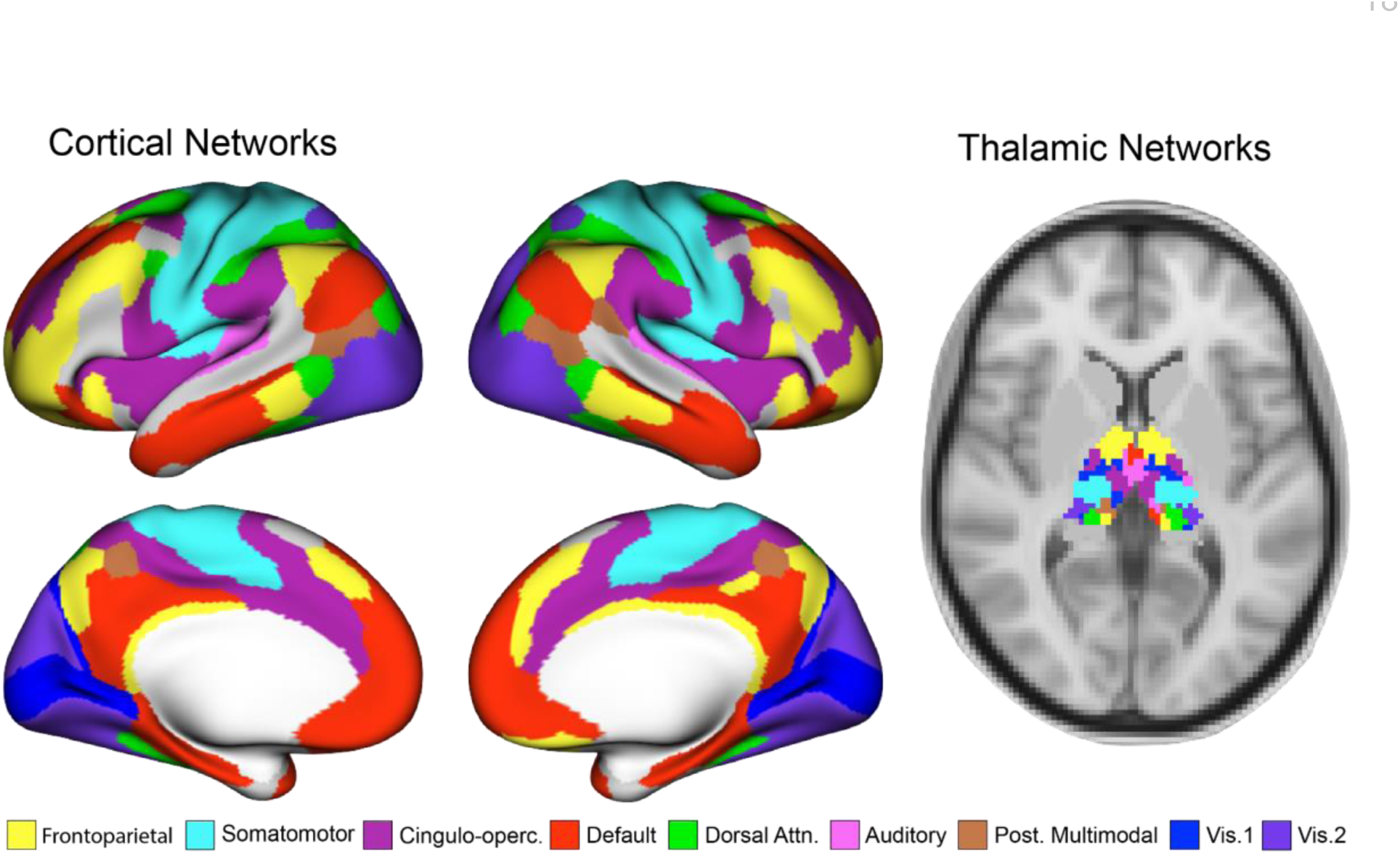
Cortical and thalamic regions for functional connectivity analysis. *Left:* nine cortical functional networks from the Cole-Anticevic Brain-wide Network Partition (CABNP) (35). *Right:* the same functional networks in the thalamus. Network thalamocortical connectivity (TCC) was computed between the mean fMRI time series in corresponding cortical and thalamic regions.

#### Modeling age trajectories

Non-linear relationships between age and TCC in 22qDel and TD cohorts were assessed with general additive mixed models (GAMMs) as in Jalbrzikowski et al., 2022 (38). Like linear mixed effects models, GAMMs can account for repeated within-subject measures with random effects. Non-linear curves are estimated with basis functions, with overfitting prevented by penalization of polynomials and restricted estimation of maximum likelihood (39–41). We examined the effects of age on TCC separately in 22qDel and TD cohorts because GAMMs allow the shape of the relationship between the smoothed predictor and dependent variable to differ between groups. For each network, a GAMM was fitted predicting TCC from the smoothed effect of age and group, controlling for sex and scanner type, with a random intercept for subject ID. Test statistics were computed for the effect of age in each group, and *p*-values were corrected for multiple comparisons with False Discovery Rate (42). Secondary analyses were performed to assess the impact of outliers, scanner type, movement, medication status, cardiac defect diagnosis, and global signal regression (GSR; see **Supplemental Methods**).

### Results

#### Non-linear age trajectories

For all nine networks in TD controls, there were no significant relationships between TCC and age after multiple comparison correction (see **Table 2**). In contrast, in 22qDel three networks (frontoparietal, somatomotor, and cingulo-opercular) exhibited a significant effect of age on TCC after FDR correction. Analysis of the 95% confidence intervals (CI) of the first derivatives of the TCC age curves identified age ranges in which significant change occurred (see **Table 2**). Specifically, frontoparietal connectivity in 22qDel increased between ages 7.5 and 12.8, relative to TD controls, while somatomotor and cingulo-opercular TCC decreased between ages 6 and 22.7. Based on the 95% CI for the group difference in age curves (see **Table 2**), 22qDel frontoparietal connectivity was significantly lower than TD in childhood, from ages 6 to 9.6, but higher in late adolescence, from age 16.8 to 19. In 22qDel, Somatomotor TCC was higher than TD prior to age 14.5, but switched to lower than TD after age 14.8. Cingulo-opercular TCC showed a similar pattern of age-related decrease in both groups, and thus did not significantly differ between groups at any age. See **Figure 3** for a visualization of frontoparietal and somatomotor age effects. Visualizations of all nine networks are included in **Figure S1**.

**Figure 3.**
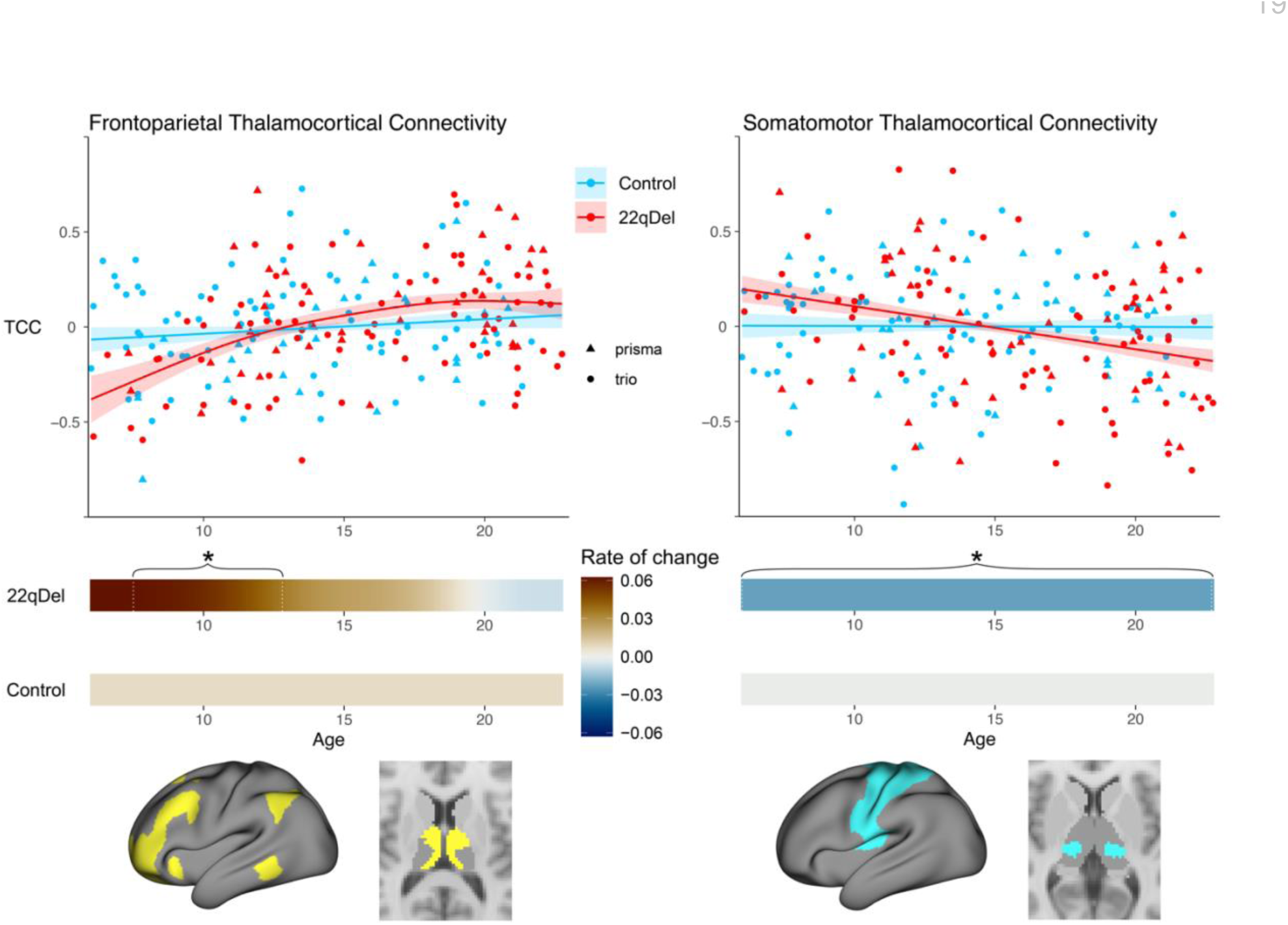
Age trajectories of frontoparietal and somatomotor thalamocortical connectivity (TCC). TCC vs age curves in 22qDel and typically developing controls. **A)** *Upper:* smoothed age curves and partial residuals for frontoparietal TCC from the GAMM predicting TCC from age, diagnosis, sex, and scanner, with a random intercept for repeated measures within subjects. The partial residual plots reflect the relationship between age and TCC, given the other covariates in the model. *Middle:* 1^st^ derivatives of the TCC vs age curve in patients and controls, with intervals of significant change determined where the 95% confidence interval for the 1^st^ derivative does not include zero, marked with brackets and asterisk. No change in frontoparietal TCC across the age range for controls, but 22qDel increases across ages 7.5-12.8. *Lower:* cortical and thalamic regions used for TCC measure. **B)** Same as **A** for the GAMM predicting somatomotor thalamocortical connectivity, showing no change across the age range in controls but a negative slope across the full age range for 22qDel.

**Table 2.**
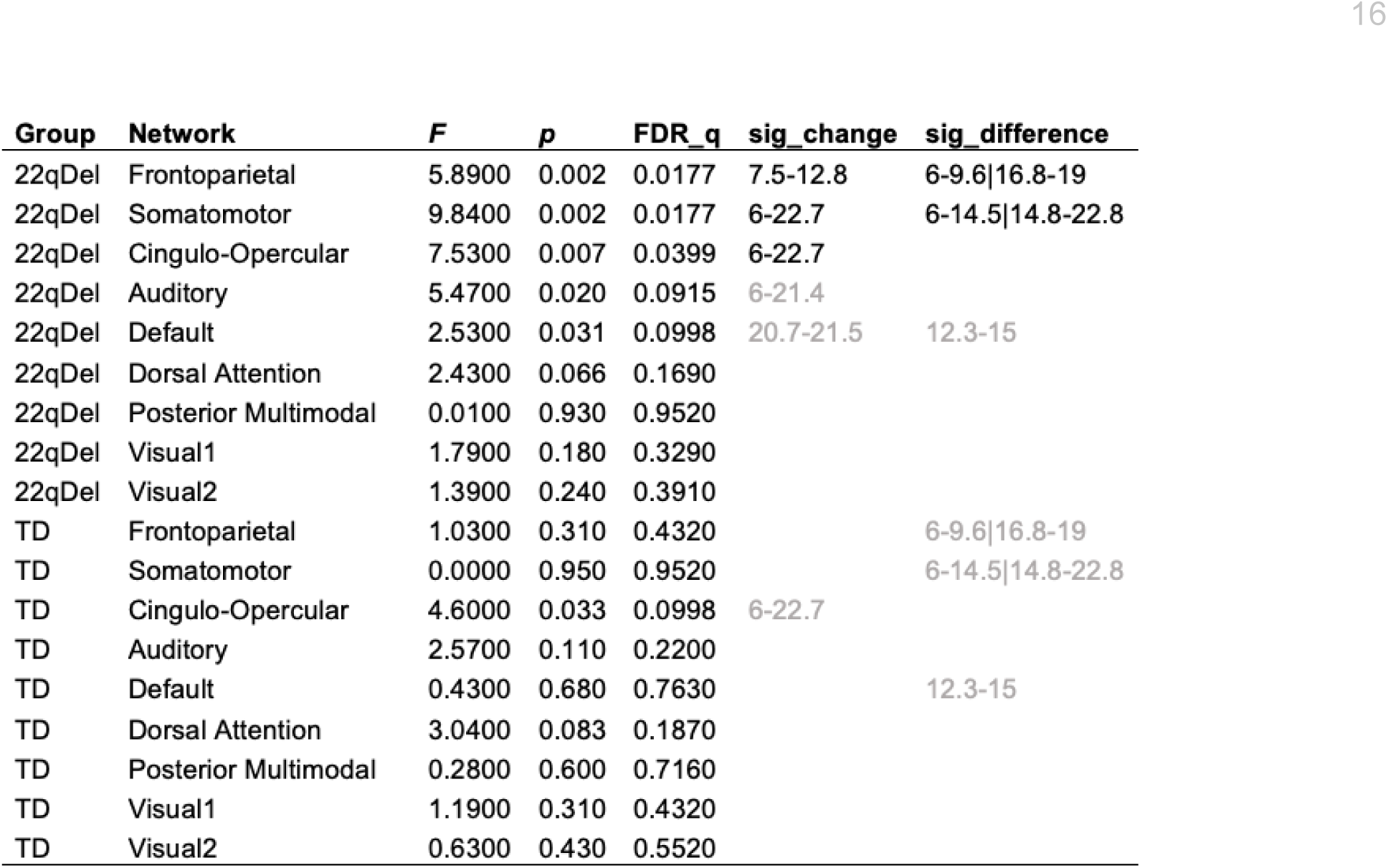
Effects of age on thalamocortical connectivity (TCC). Generalized Additive Mixed Model (GAMM) predicting TCC from age, diagnosis, sex, and scanner, with a random intercept for repeated measures within subjects. For each network, a separate TCC age curve was modeled for 22qDel and TD control groups. *F*-values and *p*-values are reported for each age effect, as well as FDR-corrected *q*-values (calculated from the set of n=18 *p*-values). “sig_change” denotes age ranges with significant change in TCC, determined where zero is not included in the 95% confidence interval (CI) for the 1^st^ derivative of the TCC vs age GAMM. “sig_difference” denotes age ranges with significant group differences in TCC, based on the 95% CI. If multiple discontinuous periods of significant difference are found for a network, they are separated by “|”. sig_change and sig_difference values are reported in grey for age curves where FDR_q>0.05.

#### Secondary analyses

Various iterations of the final model were tested to confirm robustness to potential confounds (see **Supplemental Methods**). Test statistics and probabilities are reported in the **Supplemental Results**. Conclusions from the models were not altered by inclusion of movement, medication status (antipsychotic, yes/no), or history of congenital cardiac diagnosis. Results were also robust to 90% Winsorization (i.e., restricting outliers to the 5th and 95th percentiles) prior to testing GAMMs, and the exclusion of one scanner (i.e., using only Trio data, excluding Prisma).

#### Effects of Global Signal Regression

Analyses were repeated with fMRI inputs that had not been subjected to GSR as a denoising step. Without inclusion of GSR, the pattern for somatomotor and cingulo-opercular TCC was similar but was reduced to a trend level after multiple comparison correction (see **Table S6**, **Figure S2**). The smoothed age effect on TCC no longer met multiple correction adjusted ɑ=0.05 for any network in 22qDel or TD. Despite the inclusion of motion scrubbing and nuisance regression of motion parameters, quality control functional connectivity (QC-FC) analysis showed that inclusion of GSR additionally reduced the relationship between motion and whole-brain FC in this sample, suggesting that GSR improves the data with respect to motion (see **Figure S3**).

### Discussion

Altered FC between the thalamus and brain regions involved in somatomotor and frontoparietal networks has been implicated in cross-sectional studies of individuals with schizophrenia, those at CHR for psychosis, and 22qDel (11,13,14). However, little is known about how thalamic connectivity develops with age in genetic high-risk conditions such as 22qDel. This is the first study to investigate developmental trajectories of thalamic FC in in this population. We used a powerful and flexible GAMM approach to map linear and non-linear age-related changes in network-level thalamocortical connectivity, assessed via rs-fMRI in an accelerated longitudinal cohort of individuals with 22qDel and matched TD controls, ages 6 to 23 years. This novel GAMM approach has only recently been applied for the first time to case-control neuroimaging investigations, and has identified altered developmental trajectories of structural MRI phenotypes in 22q11.2 CNVs (38).

We found that 22qDel patients exhibited significant age-related increases in frontoparietal TCC and decreases in somatomotor TCC. Frontoparietal connectivity increased steeply during childhood and the rate of change slowed during adolescence, whereas somatomotor connectivity decreased consistently through the age range. TCC was generally stable across the studied age range in TD controls. TCC in the cingulo-opercular network was also found to significantly decrease across the age range in 22qDel, while the TD group showed a similar pattern, trending towards significance in the same direction.

#### Development of thalamocortical functional connectivity in 22qDel and TD youth

These results expand on our prior findings from a cross-sectional analysis in a smaller subset of this dataset where, controlling for age, whole-thalamus FC in 22qDel relative to controls was found to be significantly increased to somatomotor regions and decreased to regions involved in the frontoparietal network (14). Our new longitudinal analysis suggests that the younger 22qDel patients were likely driving the previously observed finding of somatomotor hyper-connectivity and frontoparietal hypo-connectivity, and that this phenotype may normalize or even reverse to an abnormal extent during adolescence. Here, the bi-directional pattern of somatomotor and frontoparietal thalamocortical disruptions in 22qDel can be seen to extend to developmental trajectories. In 22qDel; frontoparietal TCC increases significantly with age, particularly prior to age 13, while somatomotor TCC decreases across the age range, intersecting with the TD curve in early/mid adolescence (see **Table 2** and **Figure 3**). Notably, GAMMs were analyzed across all nine networks represented in the thalamus, in a data driven approach, the results of which support our initial hypothesis of preferential disruptions in somatomotor and frontoparietal TCC.

Only a small number of fMRI studies have investigated typical development of thalamic FC in childhood and adolescence. A recent study in 107 TD participants found a linear decrease in salience network (cingulo-opercular) thalamic connectivity across ages 5-25 (28). Another recent study of thalamocortical FC in a large community sample (Philadelphia Neurodevelopmental Cohort), which included 1100 total scans, comprised of TD youth as well as individuals with psychosis spectrum disorders and other psychopathology, found a negative linear association between age and somatosensory thalamic connectivity, but no age by psychiatric group interactions (27). Unlike the previous two described studies, an analysis of thalamic connectivity in 52 TD individuals, which treated age as a categorical variable (child, adolescent, adult) found greater thalamic-frontal FC in adults compared to children (43). In the context of this literature, our results can be seen to generally replicate the finding of normative age-related decreases in salience network connectivity (28) in both 22qDel and TD (for whom the effect of age on cingulo-opercular TCC trended towards significance and the 95% CI of the first derivative didn’t include zero; see **Table 2**). It is still not clear how much age-related change is to be expected in typical development of somatomotor and frontoparietal thalamocortical networks (27,43). Our finding of significant age effects in these networks for 22qDel but not TD youth could be explained by either a pathological developmental mechanism in 22qDel or a compensatory “exaggeration” of typical developmental pathways. Future research in 22qDel mouse models can shed light on genetic and cellular mechanisms underlying age-related FC disruptions, which may in turn suggest potential interventions for such aberrant maturational patterns.

#### Strengths, limitations, and future directions

This study has several key strengths that support the reliability of our findings. The sample size of 112 scans from 65 22qDel patients is large for this population or similar rare disorders (44). We took advantage of an accelerated longitudinal recruitment design to map cohort-level FC-age trajectories across a key developmental window, whereas prior studies of thalamocortical FC development have relied on cross-sectional samples (27,28,43). Our GAMM analyses took advantage of this longitudinal design, with the additional benefit of being able to capture developmental trajectories whose shapes differ between cohorts (39). Potential confounds were addressed through multiple complementary approaches. To minimize the impact of scanner type, we used the longitudinal ComBat algorithm, which was specifically adapted for longitudinal neuroimaging data and represents the state-of-the-art in batch correction methods (36). Our secondary analyses showed that our primary results were robust to outliers via Winsorization, scanner effects via exclusion of data collected on one of two scanners, and to inclusion of movement parameters, antipsychotic medication status, and congenital cardiac defect diagnosis as covariates of no interest in the final model. Together, these results support a robust finding in a unique clinical population that allows for genetics-first study of phenotypes relevant to neurodevelopmental disorders.

However, certain limitations of this study must be noted. First, this dataset is not well suited for analyses of within-subject change, which might be more informative for analyses of symptom relationships over time. Similarly, this sample is not well powered for analyses of brain-behavior relationships, particularly as effect size estimates can be artificially inflated for brain-behavior relationships in small samples (45). An additional limitation is that the GAMM results differed if fMRI inputs were used that had not been subject to GSR as a preprocessing step (see **Table S6**). Without GSR, no age effects remained significant after multiple comparison correction, although 22qDel somatomotor and cingulo-opercular connectivity trended in the same direction. QC-FC analysis, developed by Power et al. to quantify the effect of participant movement on FC (46), indicated that, despite motion scrubbing and regression of motion parameters, GSR additionally reduced the relationship between movement (framewise displacement) and FC. To reduce the impact of motion on FC results in our neurodevelopmental sample, and for consistency with our prior work and other similar studies in the field (14,28), we thus present our primary results with GSR included.

Future research directions include mapping thalamocortical FC development in other clinical populations including autistic individuals, youth at CHR for psychosis, and individuals with other neuropsychiatric CNVs. Characterizing changes across the lifespan, including early through late adulthood, in these populations will also be valuable. Other methods for assessing TCC related phenotypes, such as EEG and sleep spindle detection, will be highly informative in 22qDel and related conditions (47). Additionally, the high construct validity of 22qDel animal models will allow for testing molecular and cell/circuit level hypotheses about development in 22qDel.

#### Conclusions

This study is the first to characterize longitudinal age-related changes in thalamocortical functional connectivity in children and adolescents with 22qDel. Using a novel functional atlas approach to investigate network-specific thalamocortical connectivity, we found that children with 22qDel exhibit altered maturation of these functional networks, involving a pattern of increased thalamocortical FC in the somatomotor network, concomitant with decreased connectivity in the frontoparietal network relative to TD controls. This pattern normalizes by early/mid adolescence, and potentially reverses by late adolescence. TD controls do not show the same age-related changes in frontoparietal and somatomotor connectivity. Future research in animal and *in vitro* models can shed light on biological mechanisms underlying the observed alterations in FC development in 22qDel.

## Author contributions

**Charles H. Schleifer**: Conceptualization, Methodology, Software, Formal Analysis, Data Curation, Writing - Original Draft, Writing - Review & Editing, Visualization

**Kathleen P. O’Hora**: Data Curation, Validation, Writing - Review & Editing

**Maria Jalbrzikowski**: Conceptualization, Methodology, Software, Writing - Review & Editing

**Elizabeth Bondy**: Data Curation

**Leila Kushan-Wells**: Investigation, Resources, Data Curation, Project Administration

**Amy Lin**: Data Curation

**Lucina Q. Uddin**: Conceptualization, Methodology, Writing - Review & Editing

**Carrie E. Bearden**: Conceptualization, Investigation, Writing - Review & Editing, Supervision, Project Administration, Funding Acquisition.

## Data availability

Data are publicly available from the National Institute of Mental Health Data Archives: https://nda.nih.gov/edit_collection.html?id=2414

To facilitate reproducibility and rigor, analysis code is publicly available on GitHub: https://github.com/charles-schleifer/22q_tcc_longitudinal

## Supplemental Information

### Supplemental Methods

#### Participants

The total longitudinal sample consisted of 220 scans from 135 participants (6–23 years of age; n= 65 22qDel baseline, 63.1% female; n= 69 TD controls baseline, 49.3% female), recruited from an ongoing longitudinal study at the University of California, Los Angeles (UCLA). A prior cross-sectional study of whole-thalamus FC in 22qDel included a single time point from 79 individuals included in the current longitudinal sample (14). The 22qDel participants all had a molecularly confirmed 22q11.2 deletion. 22qDel and TD participants were statistically matched based on baseline age, sex, handedness, and fMRI motion (percent frames flagged based on displacement/intensity thresholds recommended by Power et al. 2012 (31)), as well as mean number of longitudinal visits and interval between visits, using appropriate tests (ANOVA, or chi-squared). Exclusion criteria for all study participants were as follows: significant neurological or medical conditions (unrelated to 22q11.2 deletion) that might affect brain structure, history of head injury with loss of consciousness, insufficient fluency in English, and/or substance or alcohol use disorder within the past 6 months. As we aimed to include a representative cohort of CNV carriers, patients with cardiac-related issues were not excluded, as this is a hallmark of 22qDel. Healthy controls were free from significant intellectual disability and did not meet criteria for any psychiatric disorder, with the exception of attention deficit-hyperactivity disorder or a past episode of depression, due to their prevalence in childhood and adolescence (48–50). After study procedures had been fully explained, adult participants provided written consent, while participants under the age of 18 years provided written assent with the written consent of their parent or guardian. The UCLA Institutional Review Board approved all study procedures and informed consent documents.

#### Clinical assessment

At each study time point, demographic information and clinical measures were collected for each participant by trained Master’s-level clinicians, supervised by a clinical psychologist. Psychiatric diagnoses were established with the Structured Clinical Interview for DSM-IV (SCID) (51). Verbal IQ was assessed via the Wechsler Abbreviated Scale of Intelligence (WASI) Vocabulary subtest, and nonverbal IQ was assessed via the WASI Matrix Reasoning subtest. Psychiatric and dimensional psychotic-like symptoms were assessed via the Structured Interview for Psychosis-Risk Syndromes (SIPS) (52). For more details on study ascertainment and recruitment procedures, see Jalbrzikowski et al. 2012 and 2013 (53,54)

#### Neuroimaging acquisition

All subjects were imaged at the UCLA Center for Cognitive Neuroscience on either a Siemens TimTrio or Prisma scanner. The Prisma data were collected with Human Connectome Project (HCP)-style sequences. 420 volumes (5.6 min) of resting BOLD data were acquired in 72 interleaved slices with multiband-8 acceleration (voxel size = 2 × 2 × 2 mm, TR = 800 ms, TE = 37 ms, flip angle = 52°, FOV = 208 × 208 mm), along with single-band reference images and a pair of spin-echo field maps with phase encoding in the anterior-posterior (AP) and posterior-anterior (PA) directions. T1w MP-RAGE and T2w SPC images were collected in 208 sagittal slices (voxel size = 0.8 × 0.8 × 0.8 mm, FOV = 256 × 256 mm) with (T1w TR = 2400 ms, TE = 2.22 ms) and (T2w TR = 3200 ms, TE = 563 ms). The TimTrio resting BOLD data were acquired in 34 interleaved axial slices using a fast gradient-echo, echo-planar sequence (voxel size = 3 × 3 × 4 mm, TR = 2000 ms, TE = 30 ms, flip angle = 90°, FOV = 192 × 192 mm). Acquisition lasted 5.1 min and produced 152 volumes. High-resolution T1w MP-RAGE images were collected in 160 sagittal slices (voxel size = 1 × 1 × 1 mm, TR = 2300 ms, TE = 2.91 ms, flip angle = 90°, FOV = 240 × 256 mm).

#### Neuroimaging preprocessing

Structural and functional MRI data were processed using the Quantitative Neuroimaging Environment & Toolbox (32) which includes an extension of the Human Connectome Project (HCP) minimal preprocessing pipeline (33) compatible with multi-band and single-band fMRI. Additional processing of the fMRI time series included bandpass filtering, motion scrubbing for frames exceeding either a framewise displacement threshold of 0.5 mm or signal change threshold of 3 (normalized root mean square difference) proposed by Power et al. (31), and spatial smoothing (4mm Gaussian full width half maximum). Scans with >50% frames flagged for motion were excluded. To correct for spatially pervasive sources of noise including latent physiological factors and unaddressed movement, final analyses were performed on the residuals of the time series after regression of movement, mean signal from the ventricles and deep white matter, and the mean global gray matter signal (34). For a detailed description of preprocessing methods, see previous work in a subset of these data (14).

#### Functional connectivity

rs-fMRI analyses were performed using the ciftiTools package in R version 4.2.2 (55). Network thalamocortical functional connectivity (TCC) was calculated based on the Cole-Anticevic Brain-Wide Network Partition (CAB-NP) (35) which provides a whole brain cortical-subcortical extension of the HCP multimodal surface parcellation (56) derived from healthy adult resting-state fMRI. For each network, TCC was computed as the Fisher z-transformed Pearson correlation between the mean BOLD time series in the cortical portion of the network and the thalamic portion of the same network. Nine networks were investigated (frontoparietal, somatomotor, cingulo-opercular, default mode, dorsal attention, auditory, posterior multimodal, primary visual, and secondary visual), and three were excluded for lack of representation in the thalamus (orbito-affective, ventral multimodal, and language).

#### Data harmonization

To harmonize data acquired on two different scanner platforms, we applied a longitudinal implementation of the ComBat algorithm using the longComBat package in R (36,37). ComBat uses empirical Bayes methods to estimate and remove site/batch effects with increased robustness to outliers in small samples compared to general linear model approaches. ComBat was initially developed for genomics data (37), and has been subsequently adapted for neuroimaging and shown to preserve biological associations while effectively removing unwanted non-biological variation associated with site/scanner (57). The longitudinal adaptation, which uses random effects to account for within-subject repeated measures, has been shown to further increase statistical power in longitudinal neuroimaging analyses (36). LongComBat has been used to harmonize structural MRI features in a longitudinal analysis of cortical thickness and volume in a largely overlapping cohort of individuals 22qDel and controls (38).

#### Modeling age trajectories

Nonlinear relationships between age and thalamocortical connectivity in 22qDel and TD youth were assessed with general additive mixed models (GAMMs) as in Jalbrzikowski et al., 2022 (38) using the mgcv package in R. Like linear mixed effects models, GAMMs can account for repeated within-subject measures with random effects. Non-linear curves are estimated with basis functions, with overfitting prevented by penalization of polynomials and restricted estimation of maximum likelihood (39–41). We examined the smoothed effects of age on TCC separately in 22qDel and TD cohorts because GAMMs allow the shape of the relationship between the smoothed predictor and dependent variable to differ between groups. For each network, a GAMM was fitted predicting TCC from the smoothed effect of age and group, controlling for sex, scanner, and with a random intercept for subject ID. Test statistics were collected for the smoothed effect of age in each group, and *p*-values were corrected for multiple comparisons with False Discovery Rate (42). To identify age ranges of significant TCC change in each group, the first derivative of the age curve was taken, and ages in which the 95% confidence interval (CI) did not include zero for the first derivative were considered to represent significant age-associated change. Similarly, age ranges with significant group differences in TCC were determined where the 95% CI for the group difference in age smooths did not include zero.

#### Secondary analyses

To test the robustness of our models to various assumptions we performed a range of additional analyses. In our primary analyses, movement was corrected by scrubbing and by regression of motion traces from the BOLD time series, but as a supplementary analysis we repeated the main GAMMs with motion (percent frames flagged) as an additional covariate. Similarly, additional separate models were tested with antipsychotic medication status or congenital cardiac diagnosis. To test robustness to outliers, 90% Winsorization was performed, restricting extreme values to the 5th and 95th percentiles, and the main GAMM analysis was repeated on this modified input. Additionally, we repeated the main analyses with inputs that had not been subjected to global signal regression (GSR) as a preprocessing step. To quantify the relationship between movement and FC with and without GSR, we performed QC-FC analysis, an approach developed by Power et al. (46). To test the impact of using data from two scanners, the main GAMM analysis was repeated with only the Trio data (Prisma scans excluded). The Prisma cohort alone did not have enough scans to test an equivalent model.

## Supplemental Results

**Table S1.** Excluding Prisma data. GAMM age coefficients, *p*-values, FDR corrected *q*-values, and periods of significant change, tested using only data from the Siemens TimTrio scanner.

| Group | Network | F | p | FDR_q | sig_change | sig_difference |
| --- | --- | --- | --- | --- | --- | --- |
| 22qDel | Frontoparietal | 4.31 | 0.007 | 0.0375 | 9.4-12.5 | 6-10.4\|15.8-20.4 |
| 22qDel | Somatomotor | 11.97 | 0.001 | 0.0127 | 6-22.7 | 6-14.3\|14.6-22.8 |
| 22qDel | Cingulo Opercular | 8.7 | 0.004 | 0.0342 | 6-22.7 |  |
| 22qDel | Auditory | 7.15 | 0.008 | 0.0375 | 6-22.7 |  |
| 22qDel | Default | 1.72 | 0.13 | 0.282 |  | 7.2-7.9\|13.3-14.1 |
| 22qDel | Dorsal Attention | 3.03 | 0.1 | 0.262 |  |  |
| 22qDel | Posterior Multimodal | 1.16 | 0.28 | 0.371 |  |  |
| 22qDel | Visual1 | 1.32 | 0.25 | 0.371 |  |  |
| 22qDel | Visual2 | 1.06 | 0.43 | 0.486 |  |  |
| TD | Frontoparietal | 0.02 | 0.98 | 0.98 |  | 6-10.4\|15.8-20.4 |
| TD | Somatomotor | 0 | 0.98 | 0.98 |  | 6-14.3\|14.6-22.8 |
| TD | Cingulo Opercular | 3.31 | 0.031 | 0.093 | 7.3-10.1 |  |
| TD | Auditory | 1.13 | 0.29 | 0.371 |  |  |
| TD | Default | 0.63 | 0.43 | 0.486 |  | 7.2-7.9\|13.3-14.1 |
| TD | Dorsal Attention | 5.17 | 0.025 | 0.0891 | 6-22.7 |  |
| TD | Posterior Multimodal | 1.99 | 0.16 | 0.32 |  |  |
| TD | Visual1 | 1.66 | 0.22 | 0.371 |  |  |
| TD | Visual2 | 1.14 | 0.29 | 0.371 |  |  |

**Table S2.** Winsorization for outliers. Repeat of main analyses (with full TimTrio + Prisma dataset) with the additional step of 90% Winsorization, which transforms all outliers above the 95th percentile to the 95th percentile, and all below the 5th percentile to the 5th percentile.

| Group | Network | F | p | FDR_q | sig_change | sig_difference |
| --- | --- | --- | --- | --- | --- | --- |
| 22qDel | Frontoparietal | 5.96 | 0.002 | 0.0174 | 7.6-13.4 | 6-10.1\|16.3-19.5 |
| 22qDel | Somatomotor | 9.09 | 0.003 | 0.0174 | 6-22.7 |  |
| 22qDel | Cingulo Opercular | 9.48 | 0.002 | 0.0174 | 6-22.7 |  |
| 22qDel | Auditory | 4.52 | 0.035 | 0.104 | 6-22.7 |  |
| 22qDel | Default | 2.8 | 0.019 | 0.0858 | 10.4-11.5\|20.6-21.6 | 12.1-15 |
| 22qDel | Dorsal Attention | 2.38 | 0.067 | 0.151 |  |  |
| 22qDel | Posterior Multimodal | 0.04 | 0.85 | 0.846 |  |  |
| 22qDel | Visual1 | 2.59 | 0.11 | 0.197 |  |  |
| 22qDel | Visual2 | 1.22 | 0.27 | 0.442 |  |  |
| TD | Frontoparietal | 0.54 | 0.46 | 0.596 |  | 6-10.1\|16.3-19.5 |
| TD | Somatomotor | 0.04 | 0.84 | 0.846 |  |  |
| TD | Cingulo Opercular | 4.46 | 0.024 | 0.0864 | 7.8-16.4 |  |
| TD | Auditory | 3.68 | 0.057 | 0.145 |  |  |
| TD | Default | 0.56 | 0.66 | 0.744 |  | 12.1-15 |
| TD | Dorsal Attention | 2.95 | 0.088 | 0.175 |  |  |
| TD | Posterior Multimodal | 0.24 | 0.63 | 0.744 |  |  |
| TD | Visual1 | 1.04 | 0.35 | 0.488 |  |  |
| TD | Visual2 | 1.01 | 0.32 | 0.473 |  |  |

**Table S3.** Controlling for movement. Repeat of main analyses with movement (percent of frames scrubbed) as an additional fixed effect in the GAMM.

| Group | Network | F | p | FDR_q | sig_change | sig_difference |
| --- | --- | --- | --- | --- | --- | --- |
| 22qDel | Frontoparietal | 4.82 | 0.005 | 0.0331 | 7.9-12.4 | 6-9.4\|17.5-18.7 |
| 22qDel | Somatomotor | 11.11 | 0.001 | 0.0184 | 6-22.7 | 6-14.5\|14.8-22.8 |
| 22qDel | Cingulo Opercular | 9.72 | 0.002 | 0.019 | 6-22.7 |  |
| 22qDel | Auditory | 3.4 | 0.067 | 0.133 |  |  |
| 22qDel | Default | 2.38 | 0.049 | 0.111 | 20.7-21.4 | 12.3-14.8 |
| 22qDel | Dorsal Attention | 4.76 | 0.034 | 0.102 | 16.2-19.9 |  |
| 22qDel | Posterior Multimodal | 0.02 | 0.88 | 0.876 |  |  |
| 22qDel | Visual1 | 0.44 | 0.51 | 0.61 |  |  |
| 22qDel | Visual2 | 4.87 | 0.028 | 0.102 | 6-22.7 |  |
| TD | Frontoparietal | 0.65 | 0.42 | 0.585 |  |  |
| TD | Somatomotor | 0.09 | 0.76 | 0.805 |  | 6-9.4\|17.5-18.7 |
| TD | Cingulo Opercular | 6.1 | 0.014 | 0.065 | 6-22.7 | 6-14.5\|14.8-22.8 |
| TD | Auditory | 1.42 | 0.23 | 0.384 |  |  |
| TD | Default | 0.45 | 0.62 | 0.701 |  | 12.3-14.8 |
| TD | Dorsal Attention | 3.96 | 0.048 | 0.111 | 6-22.7 |  |
| TD | Posterior Multimodal | 0.49 | 0.49 | 0.61 |  |  |
| TD | Visual1 | 0.96 | 0.41 | 0.585 |  |  |
| TD | Visual2 | 2.57 | 0.11 | 0.198 |  |  |

**Table S4.** Controlling for antipsychotic medication. Repeat of main analyses with current antipsychotic medication status (yes/no) as an additional fixed effect in the GAMM.

| Group | Network | F | p | FDR_q | sig_change | sig_difference |
| --- | --- | --- | --- | --- | --- | --- |
| 22qDel | Frontoparietal | 5.8500 | 0.002 | 0.0144 | 7.5-12.8 | 6-9.6\|17-19 |
| 22qDel | Somatomotor | 10.7300 | 0.001 | 0.0144 | 6-22.7 | 6-14.5\|14.8-22.8 |
| 22qDel | Cingulo Opercular | 8.3100 | 0.004 | 0.0264 | 6-22.7 |  |
| 22qDel | Auditory | 5.2900 | 0.023 | 0.1010 | 6-22.7 |  |
| 22qDel | Default | 2.4500 | 0.046 | 0.1370 | 20.9-21.4 | 12.3-14.8 |
| 22qDel | Dorsal Attention | 3.1200 | 0.081 | 0.1950 |  |  |
| 22qDel | Posterior Multimodal | 0.0200 | 0.890 | 0.9100 |  |  |
| 22qDel | Visual1 | 2.3000 | 0.130 | 0.2360 |  |  |
| 22qDel | Visual2 | 1.3600 | 0.250 | 0.4020 |  |  |
| TD | Frontoparietal | 1.0300 | 0.310 | 0.4480 |  | 6-9.6\|17-19 |
| TD | Somatomotor | 0.0100 | 0.910 | 0.9100 |  | 6-14.5\|14.8-22.8 |
| TD | Cingulo Opercular | 4.7000 | 0.031 | 0.1130 | 6-22.7 |  |
| TD | Auditory | 2.4500 | 0.120 | 0.2360 |  |  |
| TD | Default | 0.3000 | 0.770 | 0.8620 |  | 12.3-14.8 |
| TD | Dorsal Attention | 2.9700 | 0.086 | 0.1950 |  |  |
| TD | Posterior Multimodal | 0.2600 | 0.610 | 0.7350 |  |  |
| TD | Visual1 | 1.1400 | 0.320 | 0.4480 |  |  |
| TD | Visual2 | 0.6100 | 0.430 | 0.5590 |  |  |

**Table S5.** Controlling for cardiac defect diagnosis. Repeat of main analyses with an additional covariate for lifetime congenital cardiac defect diagnosis including ventral/atrial septal defect, valvular anomalies, and conotruncal anomalies.

| Group | Network | F | p | FDR_q | sig_change | sig_difference |
| --- | --- | --- | --- | --- | --- | --- |
| 22qDel | Frontoparietal | 5.7400 | 0.002 | 0.0250 | 7.6-12.9 | 6-9.6\|17-18.7 |
| 22qDel | Somatomotor | 9.1700 | 0.003 | 0.0250 | 6-22.7 | 6-14.5\|14.8-22.8 |
| 22qDel | Cingulo Opercular | 7.1200 | 0.008 | 0.0498 | 6-22.7 |  |
| 22qDel | Auditory | 4.9400 | 0.028 | 0.1040 | 8.9-19 |  |
| 22qDel | Default | 2.5000 | 0.033 | 0.1040 | 20.8-21.4 | 12.3-15 |
| 22qDel | Dorsal Attention | 2.4800 | 0.062 | 0.1600 |  |  |
| 22qDel | Posterior Multimodal | 0.0200 | 0.900 | 0.9550 |  |  |
| 22qDel | Visual1 | 2.1000 | 0.150 | 0.2680 |  |  |
| 22qDel | Visual2 | 1.1900 | 0.280 | 0.4310 |  |  |
| TD | Frontoparietal | 1.0000 | 0.320 | 0.4420 |  | 6-9.6\|17-18.7 |
| TD | Somatomotor | 0.0000 | 0.960 | 0.9600 |  | 6-14.5\|14.8-22.8 |
| TD | Cingulo Opercular | 4.5200 | 0.035 | 0.1040 | 6-22.7 |  |
| TD | Auditory | 2.5000 | 0.120 | 0.2310 |  |  |
| TD | Default | 0.4300 | 0.680 | 0.7650 |  | 12.3-15 |
| TD | Dorsal Attention | 3.0300 | 0.083 | 0.1870 |  |  |
| TD | Posterior Multimodal | 0.2800 | 0.600 | 0.7190 |  |  |
| TD | Visual1 | 1.2700 | 0.290 | 0.4310 |  |  |
| TD | Visual2 | 0.6000 | 0.440 | 0.5630 |  |  |

**Table S6.** Omission of global signal regression (GSR). Repeat of main analyses without GSR as a preprocessing step. Age effects in 22qDel were significant at p<0.05 uncorrected for somatomotor, cingulo-opercular, auditory, and dorsal attention networks, but reduced to trend level after FDR correction.

| Group | Network | F | p | FDR_q | sig_change | sig_difference |
| --- | --- | --- | --- | --- | --- | --- |
| 22qDel | Frontoparietal | 0.63 | 0.49 | 0.617 |  |  |
| 22qDel | Somatomotor | 4.27 | 0.04 | 0.18 | 6-22.7 | 6-22.8 |
| 22qDel | Cingulo Opercular | 8.48 | 0.004 | 0.0737 | 6-22.7 |  |
| 22qDel | Auditory | 3.38 | 0.022 | 0.131 | 8.8-12.5 | 6-11.1\|16.2-22.6 |
| 22qDel | Default | 0.01 | 0.91 | 0.912 |  |  |
| 22qDel | Dorsal Attention | 3.13 | 0.018 | 0.131 | 19.3-21.5 | 20.7-22.8 |
| 22qDel | Posterior Multimodal | 0.77 | 0.38 | 0.571 |  |  |
| 22qDel | Visual1 | 1.15 | 0.28 | 0.527 |  |  |
| 22qDel | Visual2 | 2.78 | 0.097 | 0.291 |  |  |
| TD | Frontoparietal | 0.36 | 0.55 | 0.62 |  |  |
| TD | Somatomotor | 3.41 | 0.066 | 0.239 |  | 6-22.8 |
| TD | Cingulo Opercular | 1.11 | 0.29 | 0.527 |  |  |
| TD | Auditory | 1.43 | 0.23 | 0.526 |  | 6-11.1\|16.2-22.6 |
| TD | Default | 0.43 | 0.51 | 0.617 |  |  |
| TD | Dorsal Attention | 0.14 | 0.71 | 0.749 |  | 20.7-22.8 |
| TD | Posterior Multimodal | 1.56 | 0.21 | 0.526 |  |  |
| TD | Visual1 | 0.62 | 0.43 | 0.598 |  |  |
| TD | Visual2 | 0.94 | 0.33 | 0.544 |  |  |

**Figure S1.**
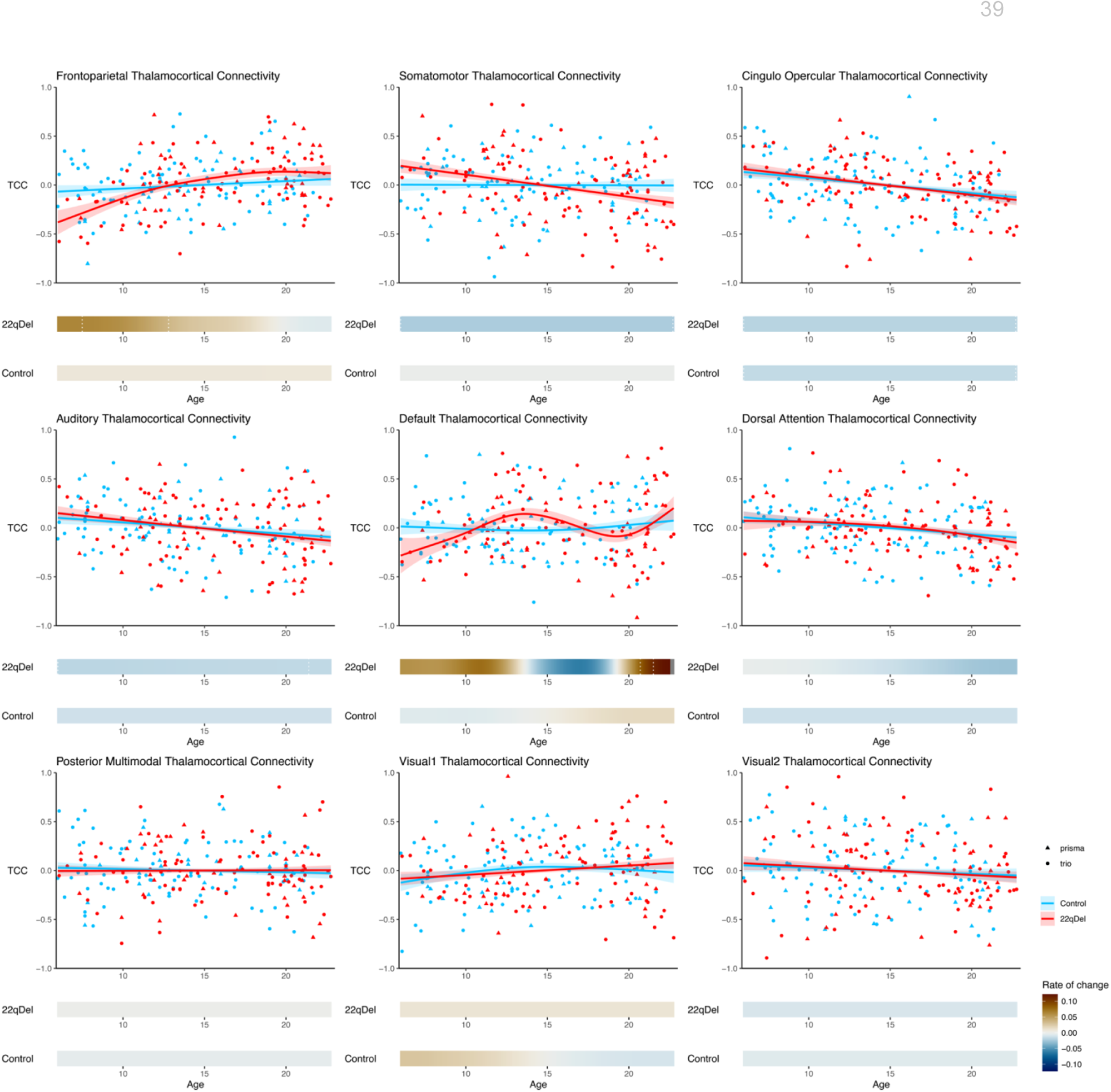
Age trajectories for all networks. Visualization of GAMM curves and partial residuals *(above)* and first derivatives *(below*) for all nine networks from the primary analysis.

**Figure S2.**
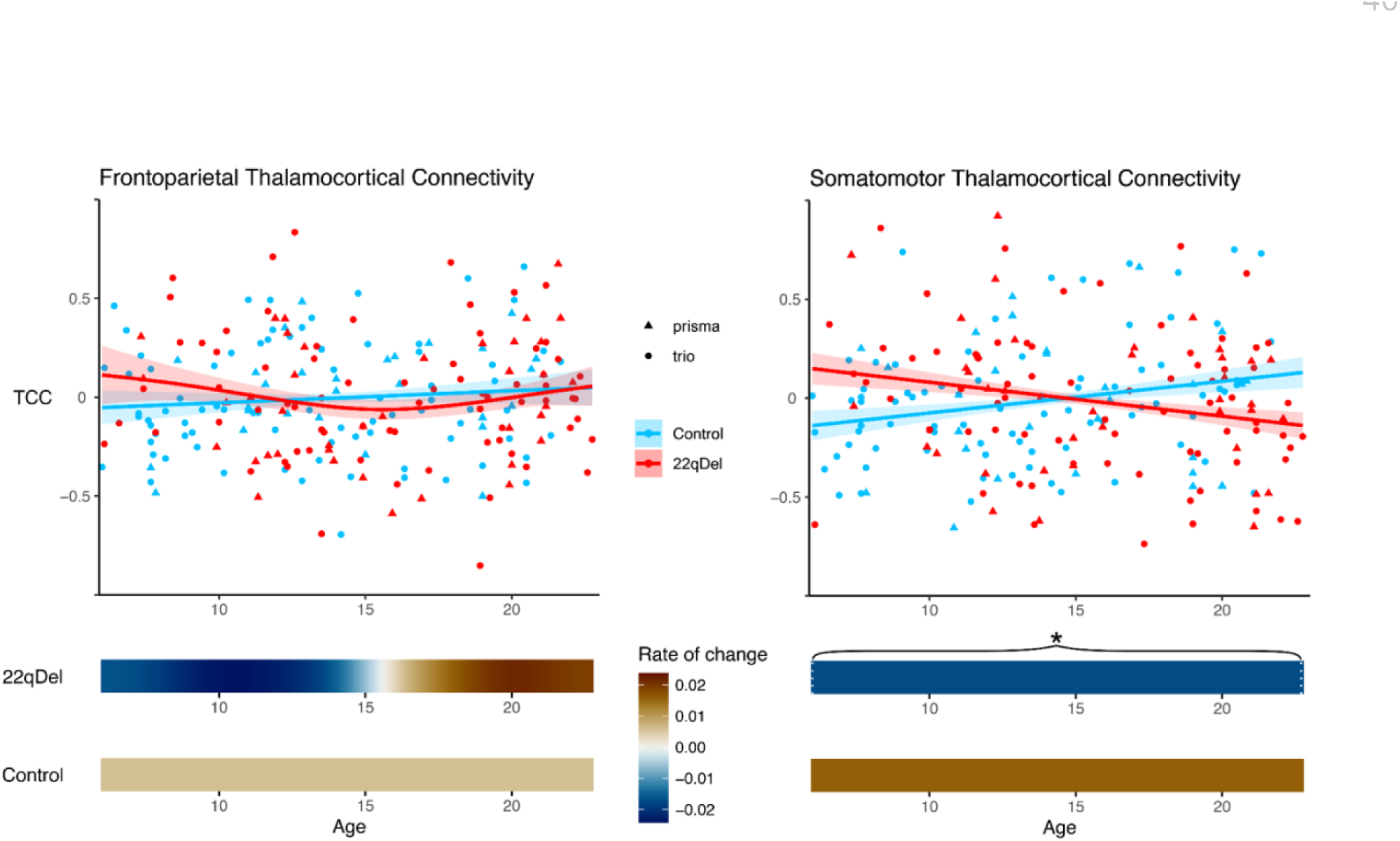
Age trajectories of frontoparietal and somatomotor thalamocortical connectivity without GSR. Visualization of GAMM curves and partial residuals *(above)* and first derivatives *(below*) for frontoparietal *(left)* and somatomotor *(right)* networks.

**Figure S3.**
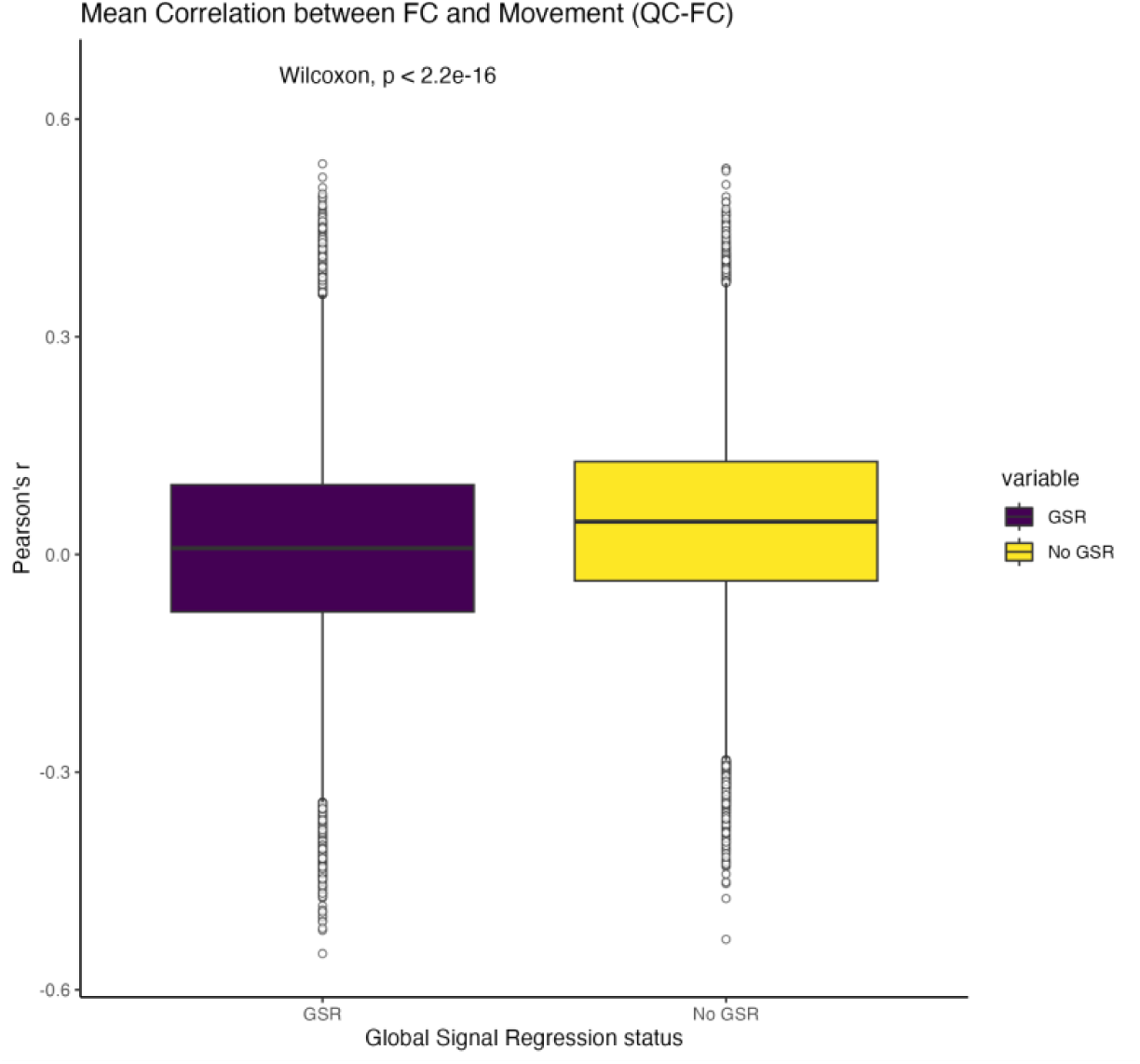
QC-FC relationships with and without GSR. For each pair of regions in the n=718 region CAB-NP parcellation, QC-FC was calculated as the Pearson correlation between functional connectivity and movement across all baseline 22qDel and TD scans (46). QC-FC values were compared for data with and without GSR using a Wilcoxon signed rank test indicating a significantly decreased relationship between movement and functional connectivity in the global signal regressed data (purple) compared to the data without GSR (yellow).

